# Biophysical properties of material-colonizing fungal biofilms: multiscale structural and mechanical characterization

**DOI:** 10.64898/2026.04.22.720134

**Authors:** Clémentine Ferrari, Abolfazl Dehkohneh, Julia Schumacher, Yu Ogawa, Ruben Gerrits, Peter Fratzl, Anna A. Gorbushina, Cécile M. Bidan

## Abstract

Black extremotolerant fungi form persistent biofilms on a wide range of natural and engineered substrates. Able to weather minerals, affect stone monument surfaces, and colonize solar panels, they demonstrate a strong capacity to interact with and modify material surfaces, even under most extreme conditions. In this study, we establish a methodological workflow for the structural and mechanical characterization of melanized biofilms formed by the black fungus *Knufia petricola*. This species represents a broader group of resilient surface colonizers and provides a model for in-depth investigation. When grown on solid agar/air interface, this species predominantly forms a compact biofilm, composed of spherical cells, while retaining the capacity for filamentous growth, providing a suitable framework to explore morphology-dependent biomechanical responses.

The proposed toolbox combines complementary analytical techniques spanning multiple spatial scales, including shear-rheology to quantify bulk viscoelastic behavior, micro-indentation to resolve local stiffness of the biofilm surface, micro-computed tomography for non-destructive three-dimensional visualization of biofilm architecture, and cryogenic preparation methods and electron microscopy for high-resolution ultrastructural analysis. As a case study, we applied this workflow to compare biofilms grown on two nitrogen sources (NO_3_^-^ vs. NH_4_^+^).

Our results reveal that the nitrogen source plays a key role in biofilm morphology across multiple hierarchical levels - ranging from cell division patterns and distribution of extracellular polymeric substances (EPS) to overall mechanical properties, where NO_3_^-^ leads to budding-dominated growth and increased stiffness, whereas NH_4_^+^ promotes meristematic growth and softer biofilms. The successful transfer and integration of methods originally developed for bacterial biofilm research highlights the feasibility of quantitative mechanical analyses in fungal systems. This multiscale toolbox provides a foundation for advancing the mechanistic understanding of fungal biofilms and biofilm-material interactions, with implications for geomicrobiology, material biodeterioration, and the design of bio-inspired functional materials.

**Graphical Abstract:** 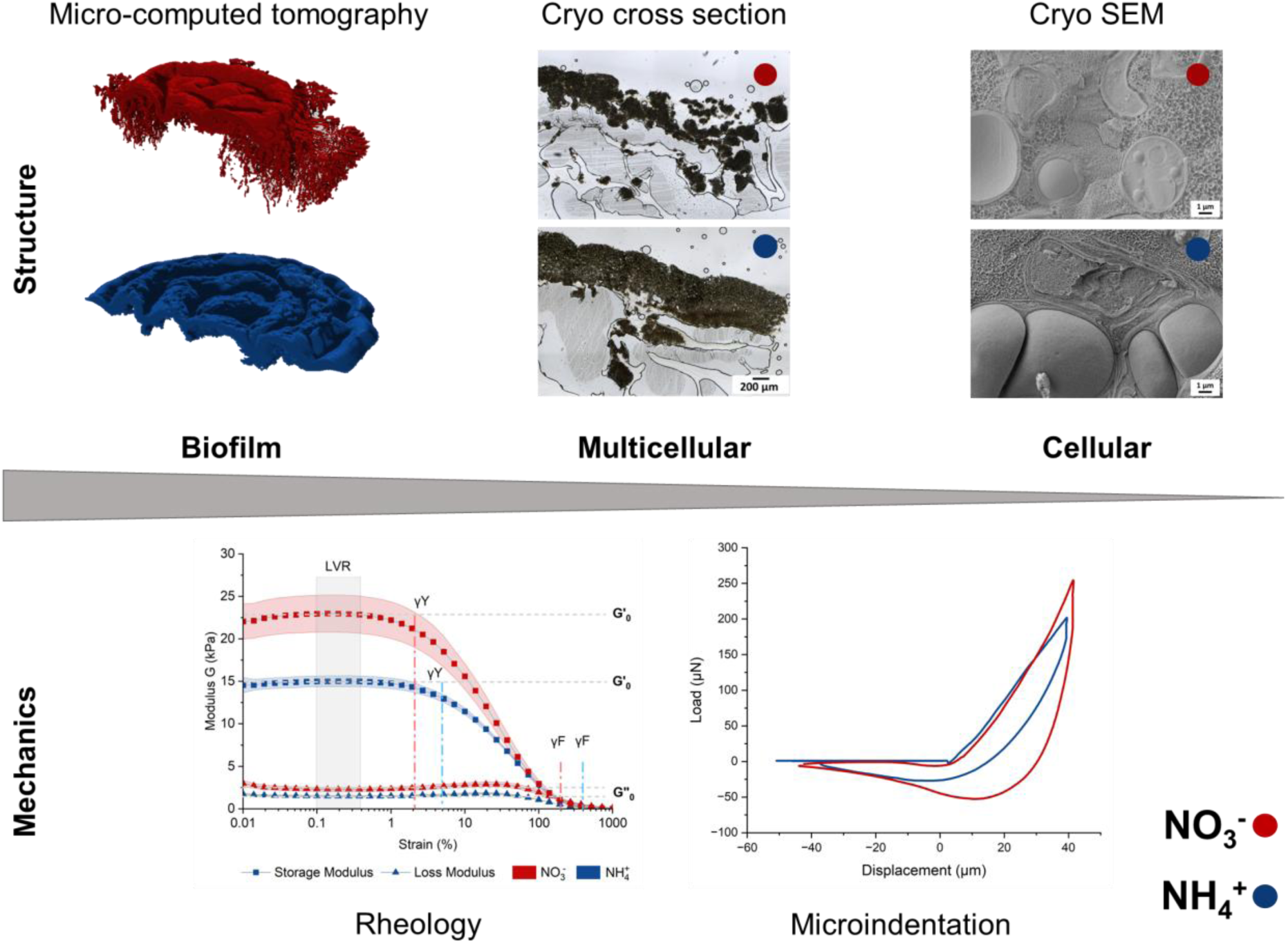

## Introduction

Biofilms are surface-attached microbial communities embedded in self-produced extracellular polymeric substances (EPS) that protect the microorganisms from biological, chemical and physical challenges as well as serve as extracellular reservoirs for nutrients. ^1, 2^ Importantly, as inherently surface-and material-bound systems, biofilms-especially bacterial biofilms-have recently gained increased attention in interdisciplinary research at the interface of microbiology, biophysics, and materials science with the aim of understanding and manipulating their structural and functional properties for applications in antimicrobial strategies and the development of engineered living materials. ^3–5^ So far, much of this work has focused predominantly on prokaryotic biofilms. In natural environments, biofilms are complex, multispecies communities comprising bacteria, archaea, fungi, and other microorganisms, whose interactions critically influence their emergent properties.

This effort has produced a sophisticated methodological toolkit for probing bacterial biofilm structure and mechanics across scales. ^6^ For example, cross-sections of *Escherichia coli* biofilms stained with a marker for the EPS were prepared in cryogenic conditions and observed by fluorescence confocal scanning microscopy to reveal typical layered structures as well as the orientation of the fibrous EPS. ^7, 8^ Cryogenic electron microscopy was used to visualize details of cellulose secretion sites in several bacterial species and revealed characteristics specific to species producing crystalline cellulose. More recently, micro-computed tomography was applied to biofilm research with the help of contrast agents to simultaneously reveal both the macroscopic morphology and the internal structure without cross-sections. ^9^

Characterizing biofilm mechanical properties has been the focus of many studies assessing biofilm adaptation to environmental influences, or the structure-function relationships in biofilms. ^4, 5^ As such, shear-rheology, indentation-based approaches, as well as methods routinely used to characterize the mechanical properties of mammalian tissues, have been adapted to biofilm systems. ^3, 6^ These techniques have revealed the key role of the EPS components and their mutual interactions in the mechanical behavior of microbial biofilms formed by various bacterial species. ^10–12^

In contrast, fungal biofilms have received far less attention despite their ecological relevance and their diverse architecture, arising entirely from growth-driven processes such as budding, hyphal extension, branching and matrix deposition, as they are non-motile. ^13, 14^ Across fungal lineages, cell walls are composed of glucans and chitin, while their thickness, outer layers and extracellular exopolysaccharides, glycoproteins, and secondary metabolites such as 1,8dihydroxynaphthalene (DHN) melanin vary with species, morphology and environmental conditions. ^15, 16^ Fungal growth forms range from unicellular budding yeasts to filamentous hyphal networks that generate structurally complex biofilms and reproduction structures. Many species exhibit pronounced morphological plasticity in response to environmental conditions, leading to substantial variations in colony architecture and biomass organization. ^15, 17^

Black fungi, however, display a unique combination of traits enabling them to grow and persist in environments where most other microbes cannot survive: slow growth, constitutive DHN melanin synthesis, absence of reproduction structures, and flexible, unconventional cell division programs, ranging from yeast-like budding to meristematic growth that generates densely packed cells with thick, layered walls. ^18, 19^ Two practical subtypes are often distinguished: black yeasts, which predominantly produce single cells and form dark, slimy colonies, and microcolonial black fungi (rock-inhabiting fungi), which form cell chains and clusters connected by their melanized cell walls and EPS matrices, resulting in small but numerous and adherent colonies that formed a base of subaerial biofilms on rocks and exposed human-made materials such as monuments, façades, and photovoltaic modules. ^20–24^ These features raise the central methodological question: can tools originally developed for bacterial biofilm analysis reliably characterize the structurally distinct architectures of black fungal biofilms? *Knufia petricola*, a representative microcolonial black fungus colonizing rocks and human-made materials, ^25–27^ provides an excellent test system for the proposed methodological transfer. Together with genetic engineering tools, the robust, non-pathogenic growth of *K. petricola* supports reproducible quantitative assays and controlled mineral-weathering experiments. ^28–31^ Several traits – including deposition of DHN melanin in the outer cell wall, accumulation of carotenoids in cell membranes, and an EPS matrix dominated by pullulan likely contribute to the development, cohesion and structural characteristics of its biofilms. ^32–34^ Moreover, its compatibility with phototrophic partners allows the generation of synthetic biofilms with tunable community composition, further expanding its versatility as a model for fungal biofilm biology. ^16, 28^ A recent study showed that the macroscale morphology of *K. petricola* biofilms differs when grown on solid media containing either nitrate (NO_3_^-^) or ammonium (NH_4_^+^) as sole nitrogen source, suggesting that environmental conditions modulate biofilm architecture.^35^ Whether these morphological differences correspond to measurable biomechanical differences remains unknown, partly because standard tools for such measurements have not yet been validated for black fungal biofilms.

In this study, we adapt and evaluate a cross-scale analytical toolbox originally developed for bacterial biofilm research and apply it to black fungal biofilms. Using two nitrogen-conditioned *K. petricola* biofilms (NO_3_^-^ vs. NH_4_^+^), we assess: [i] biofilm-scale morphology, quantified by micro-computed tomography (micro-CT); [ii] multicellular organization, visualized by cryo-section imaging; [iii] cellular-scale architecture, resolved through cryo-scanning electron microscopy (cryo-SEM); [iv] bulk mechanical behavior, probed by rheology; and [v] local stiffness at the multicellular scale, measured via micro-indentation (Figure 1).

**Figure 1:**
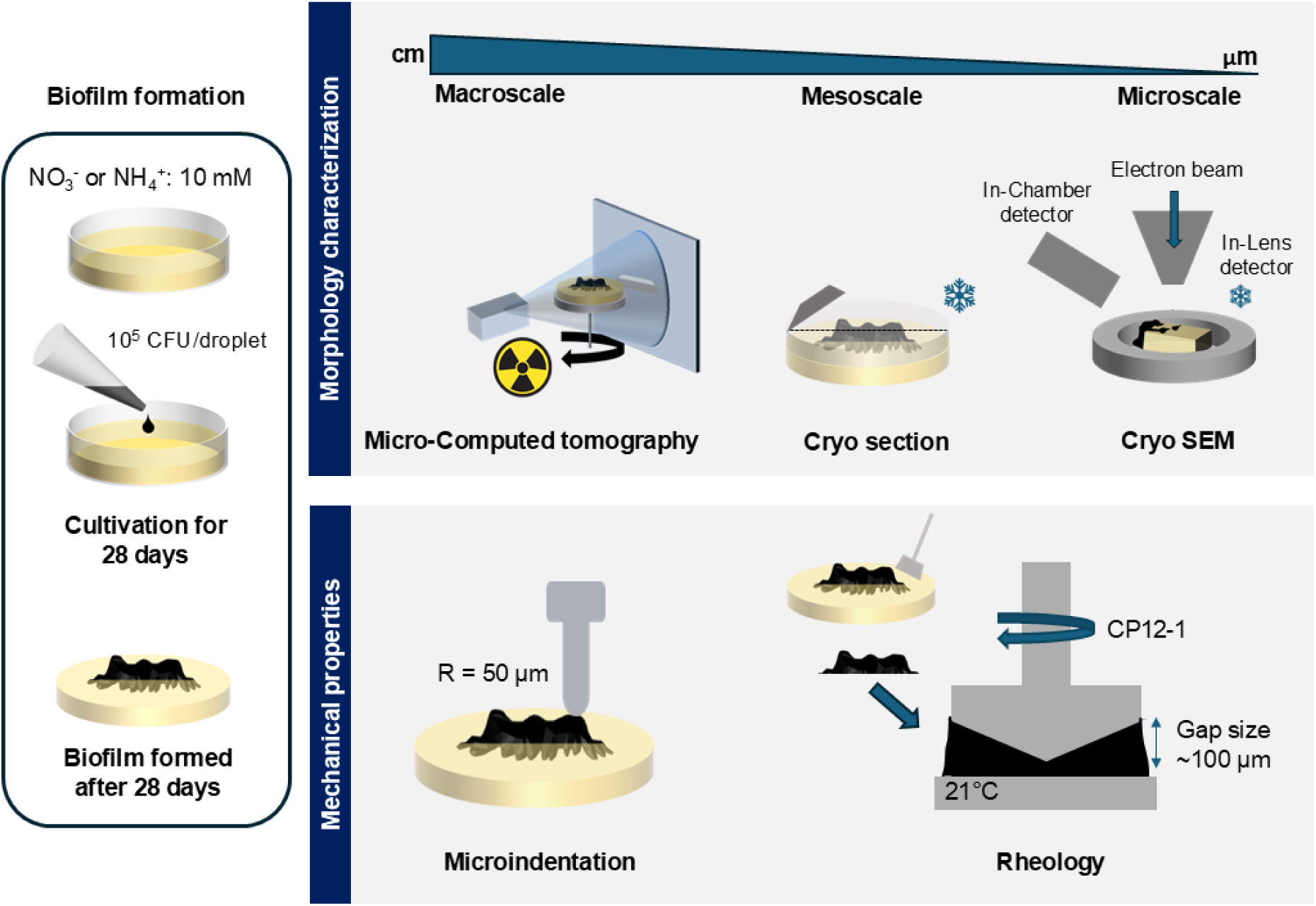
Experimental concept of multiscale morphological and mechanical characterization of *Knufia petricola* biofilms formed on agar media containing two different nitrogen sources.

Our objectives were [i] to evaluate whether methodologies developed for bacterial biofilms can be reliably transferred to black fungal biofilms, [ii] to identify scale-dependent readouts that distinguish the two condition-specific *K. petricola* biofilms, and [iii] to establish a validated workflow that enables future hypothesis-driven studies of black fungal biofilm architecture and mechanics.

## Results

The nitrogen source determines the morphology of *K. petricola* biofilms: the biofilms exhibited more filamentation at the biofilm edge and more wrinkles on the biofilm surface in the presence of NO_3_^-^, while biofilm surface was smoother and filamentation occurred under the central rings in NH_4_^+^-containing medium, as was previously observed by Dehkohneh and co-workers. ^35^ The biofilms grown on solid media containing either NO_3_^-^ or NH_4_^+^ as sole nitrogen source were tested after 28 days incubation at 25°C in darkness.

### Micro-CT reveals interactions of the biofilms with the agar substrates

To assess the morphological differences between *K. petricola* biofilms grown on media containing either NO_3_^-^ or NH_4_^+^, the native biofilms were scanned on their agar substrate by X-ray micro-CT after being stained with iodine to enhance the contrast. The three-dimensional image reconstructions revealed that the biofilms grown on NO_3_^-^-containing media were comparatively thinner than biofilms on media containing NH_4_^+^ (Figure 2A). However, increased filamentation at the edges of biofilms, along with deeper penetration into the agar, was observed in the presence of NO_3_^-^. To quantify these differences, the segmented volume of each biofilm, the distribution of their local thickness (Figure 2B) and the surface interacting with the air and with the agar substrate were measured (Figure 2C). While conserving similar volumes between 30 and 35 mm^3^ (Figure S1), biofilms grown with the two different nitrogen sources displayed distinct thickness distributions. Biofilms grown on NO_3_^-^-containing medium showed rather homogenous thickness distributions, where the upper values corresponded to the thickness of the biofilm above the agar substrate (> 150 µm) and the lower values corresponded to the bundles of filaments penetrating the agar (∼90 µm). In contrast, these two characteristic thicknesses appear to be more distinct in biofilms grown on NH_4_^+^-containing medium, as the biofilms have less filaments and a more homogeneous thickness (∼320 µm) than in biofilms grown on NO_3_^-^-containing medium. Yet, the few existing filaments are also 90 µm thick.

**Figure 2.**
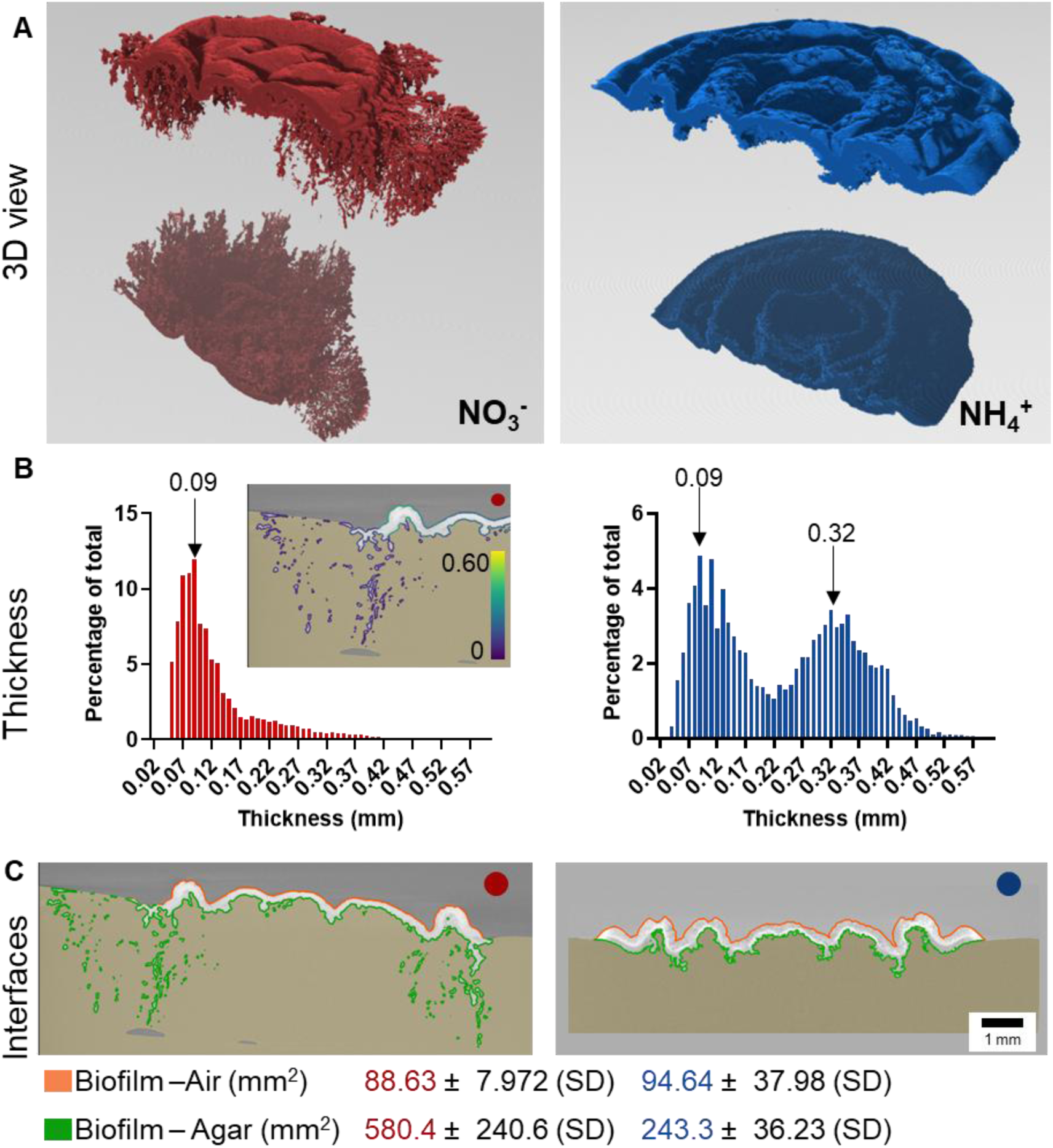
Micro-CT illustrates biofilm morphology and surface roughness in three dimensions. The red color represents the biofilms grown on NO_3_^-^ medium, and the blue color represents biofilms grown on NH_4_^+^ medium. **(A)** 3D view with a mirror effect used to simultaneously display the top and bottom of the biofilm. **(B)** Percentage of biofilm thickness distribution. Thicknesses are calculated as the diameter of a hypothetical sphere that fits within each boundary point. **(C)** Surfaces of the biofilm-air and biofilm-agar interfaces. Three biological replicates for each condition were imaged with the same voxel sizes.

To assess how much these morphological differences may impact biofilm interactions with their direct environment, the area of interface between [i] the biofilm and the air and [ii] the biofilm and the agar substrate were computed in each condition (Figure 2C). The source of nitrogen did not seem to impact the surface of exchange between the biofilm and the air (∼90 mm²). However, NO_3_^-^-containing medium promoted interactions between fungal biofilm and the substrate as their interface was more than doubled (580.4 ± 240.6 mm² for NO_3_^-^ vs. 243.3 ± 36.2 mm² for NH_4_^+^). This large interaction potential can be attributed to the thin and numerous filaments penetrating the agar (Figure 2A).

### Cryo microtome cross sections show the penetration of *K. petricola* cells in the agar

To study cell arrangement, biofilm microstructure, and interaction with the agar at the cellular level, cryosections of 30 µm-thick biofilm slices were produced and imaged in both the hydrated and dry states (Figure 3). The hydrated sections not only revealed differences in biofilm thickness between NO_3_^-^-and NH₄⁺-containing media, but also in biofilm density (Figure 3A). Indeed, NH₄⁺-containing medium seems to yield thicker biofilms than NO_3_^-^-rich medium (600 µm vs. 400 µm, respectively), and the thickness seems to be more homogeneous along the surface of the substrate for the former. In contrast, biofilms grown on NO_3_^-^-containing medium appear to be arranged in denser, smaller vertical packages with a higher cell density as indicated by the darker contrast (Figure 3A, B). Moreover, differences in cell sizes can be observed in biofilms grown on NH_4_^+^-containing medium, where the upper layer (∼400 µm) contains cells of ∼15 µm diameter, whereas lower layer closer to the substrate (∼150 µm) contains cells of ∼5 µm.

**Figure 3.**
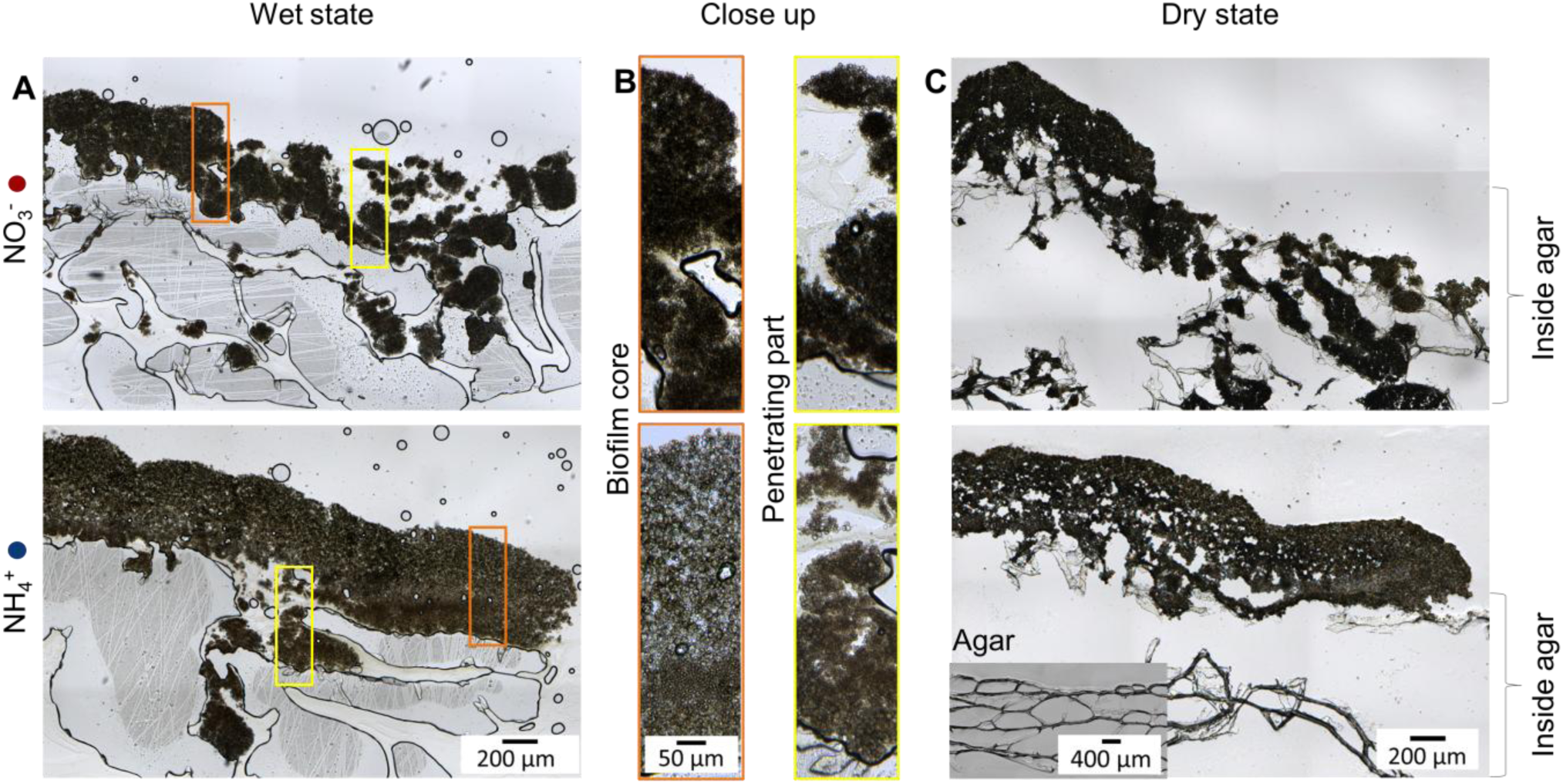
Slices of 30 µm biofilm grown on NO_3_^-^-and NH_4_^+^-containing media cut in cryogenic condition and imaged in wet **(A, B)** and dry state **(C)**. Braces on the right correspond to the part in the agar.

To better observe the interactions with the agar, the slices were allowed to dry before further imaging (Figure 3C). Biofilm filaments appeared as cell aggregates attached to the agar network, with cells penetrating deeper in NO_3_^-^-containing agar substrate. While the agar may seem disrupted in the presence of the fungal biofilm (Figure 3C inset), the effect of the sectioning process cannot be excluded. Drying the samples also shows that the homogenous biofilms grown on NH_4_^+^-containing medium have become porous, indicating that water was significantly contributing to the low density of the biofilm.

### Cryo-SEM allows to see interactions between cells and their surrounding environment

Cryo-SEM images of freeze-fractured biofilms were taken to visualize individual cells and their EPS, as well as their interactions while penetrating the agar. Two different network sizes were observed: the agar network had pores around 1 µm in size, while the EPS network had pores around tens of nanometers (Figure 4A). The interaction between the two networks (Figures 4B, C) was characterized by the EPS network integrating into the pores of the agar network in both conditions. Figure 4B shows cells surrounded by a biofilm matrix between two agar networks, which could be part of a filament penetrating the NO_3_^-^-containing agar. In biofilms grown on NH_4_^+^-containing medium, the cells tended to be more isolated from each other. However, they still showed an EPS layer on the outer cell wall (Figure 4C).

**Figure 4.**
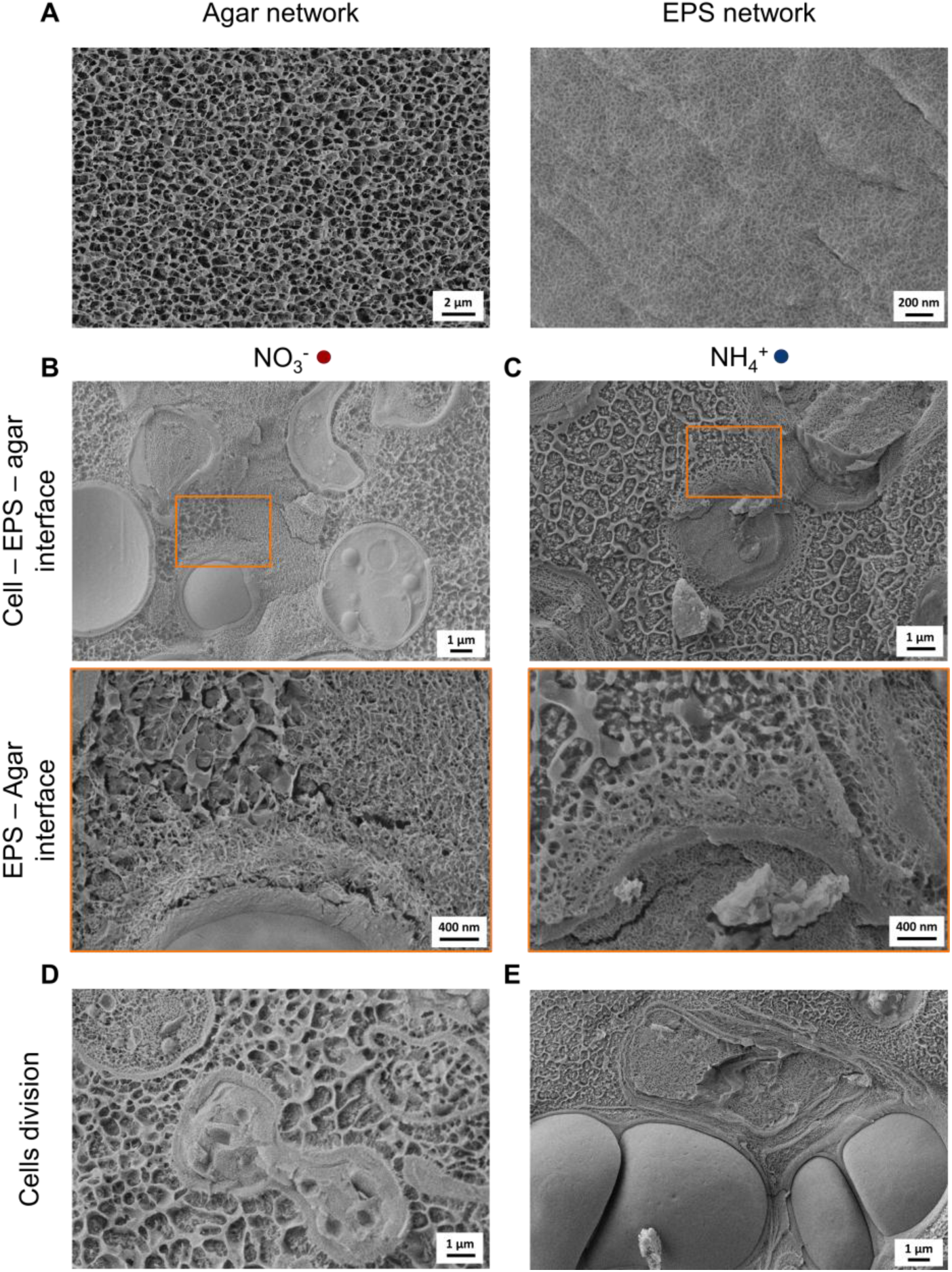
Cryo-SEM images of freeze-fractured biofilms. **(A)** Agar and EPS network isolated from the biofilm sample. **(B,C)** The interface between cells, EPS and agar. **(D,E)** Representative cell division patterns observed under different nitrogen regimes: predominantly budding in the presence of NO_3_^-^ (D) and meristematic growth in the presence of NH_4_^+^ (E).

At the cellular level, cells grown in NO_3_^-^-containing medium tended to divide frequently by budding (Figure 4D), whereas NH_4_^+^ appeared to promote meristematic growth. The latter was characterized by the formation of a cell wall between the daughter cells, whilst both were still surrounded by an old cell wall, EPS being present between the old and new walls (Figure 4E).

### Shear-rheology reveals the viscoelastic properties of *K. petricola* biofilms

To assess the mechanical consequences of the morphological and structural differences of *K. petricola* biofilms formed with different nitrogen sources, their viscoelastic properties were probed using bulk oscillatory shear-rheology (Figure 5). For this, a piece of biofilm was first scrapped off from the agar surface, so that only the core part of the biofilm was tested (not the filaments) (Figure 5A). Approximately one quarter of the biofilm was enough to fill a 100 µm gap in the cone-plate geometry of the rheometer. Amplitude sweeps were performed to determine the viscoelastic response of the fungal material to the applied shear strain (Figure 5B). For both nitrogen sources, the linear viscoelastic region (LVR) ranged from 0.1-0.268% strain, with the storage modulus (G′_0_) approximately one order of magnitude greater than the loss modulus (G″_0_), confirming the viscoelastic nature of *K. petricola* biofilms (Figure 5C, D). The significant differences between the mean values of G′_0_ and G″_0_ for the different growth conditions revealed that biofilms grown in the presence of NO_3_^-^ were about 50% stiffer than those grown on NH_4_^+^-containing medium, with storage moduli of ∼24 kPa and ∼15 kPa, respectively.

**Figure 5.**
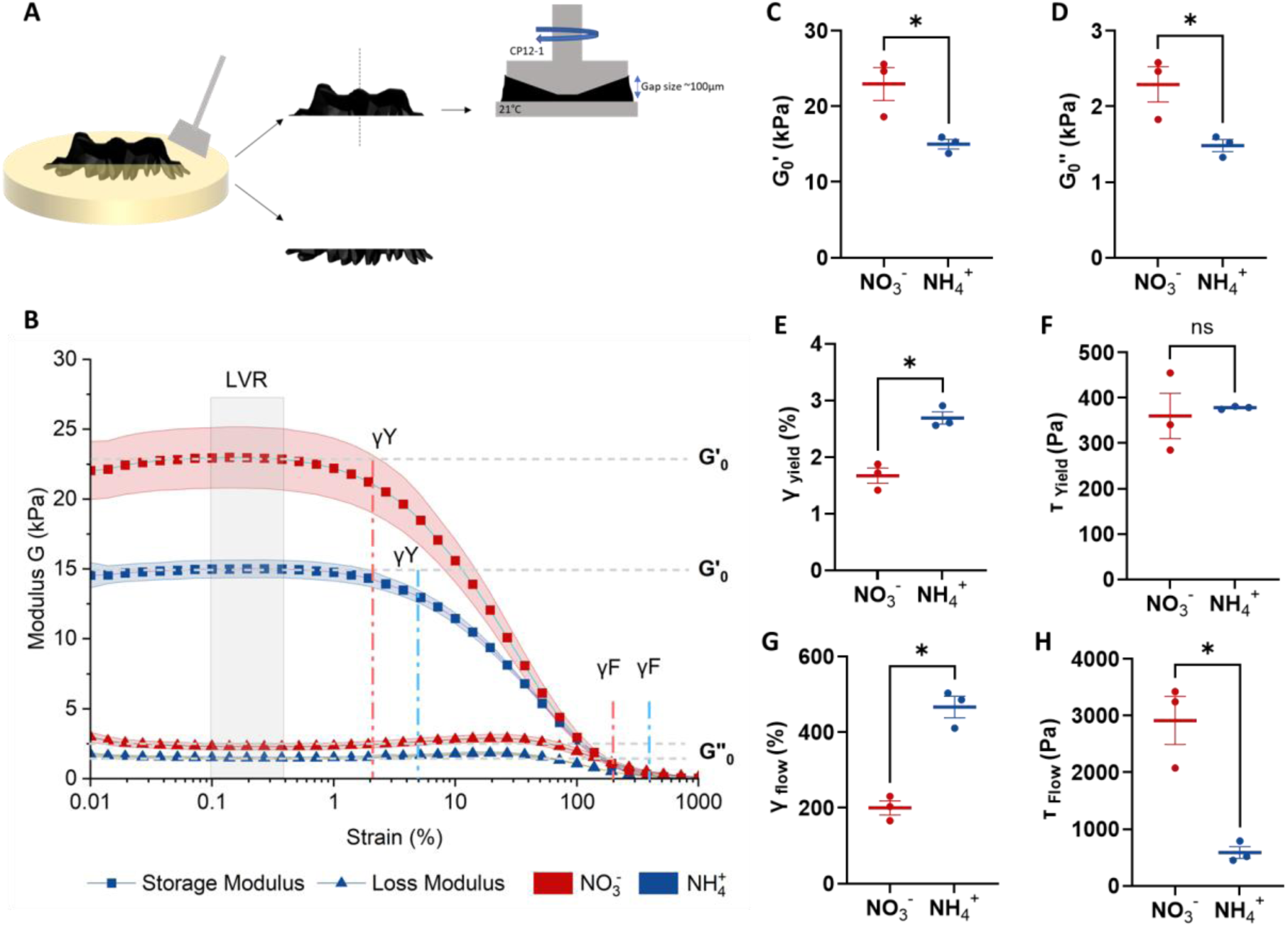
Oscillatory shear-rheology of *K. petricola* biofilms grown on media with different nitrogen sources. **(A)** Preparation of the samples. **(B)** Average strain amplitude sweeps. The linear viscoelastic region (LVR) ranged from 0.1-0.268% strain for both nitrogen sources. The γY was defined as 95% of the LVR limit and γF was determined as the strain at which the storage modulus (G′) intersected the loss modulus (G″). **(C)** Storage modulus (G′_0_). **(D)** Loss modulus (G**″_0_**). **(E)** Yield strain (γY) determined where G′**_0_** < 0.95*G′_0_. **(F)** Yield stress (τY). **(G)** Flow strain (γF). **(H)** Flow stress (τF). Statistical analysis was done with three biological replicates represented with dots on the graphs and a bar for the mean value with the standard error. A paired two tailed t-test was applied with statistical significance as p < 0.05 shown as * and p > 0.05 as ns = non-significant.

The fungal biofilms started to yield under lower shear strain when grown with NO_3_^-^compared to NH_4_^+^ (∼1.8% vs. ∼2.8% respectively), indicating that NO_3_^-^ favored the formation of biofilms that were more prone to irreversible deformations (Figure 5E). Yet, the yield stress (τ_Y_) determined as the product of the yield strain (γ_Y_) and the shear modulus *G* = √(*G*′^2^ + *G*″^2^) (Figure S2) showed no difference between the two types of samples (∼375 kPa) (Figure 5F). Similarly, biofilms grown on NO_3_^-^-containing substrates start to flow – i.e. be dominated by a viscous behavior – under lower shear strains compared to NH_4_^+^ (Figure 5G), while this order is reversed when considering the flow stress, which accounts for the difference of shear moduli between the two conditions (Figure 5H).

Biofilms grown in the presence of NH_4_^+^ were thus softer and deformed irreversibly (yield) under the same shear stress compared to biofilms grown with NO_3_^-^. However, the stress range defining the transition from material yield to flow was larger in biofilms grown with NO_3_^-^, indicating that their failure under increasing stress was more progressive compared to NH_4_^+^ (Figure S2).

### Micro-indentation allows local mechanical assessment of *K. petricola* biofilms

To evaluate the mechanical properties of *K. petricola* biofilms on a multicellular scale, indentation measurements were performed while preserving the native state and architecture of the samples. Micro-indentation with a 50 µm conospherical tip allowed measurement of multiple cells of 5 to 10 µm diameter and their EPS (Figure 6A). After completing the loading phase and reaching the maximum penetration depth set at ∼40 µm, the tip was held in position for 5 seconds (Figure 6B). During this holding time, the characteristic relaxation time was calculated by fitting an exponential decay function on the Load-Time curve. For both nitrogen sources, a relaxation time of 1.25 seconds was observed for the agar and the biofilms suggesting that similar deformation and reorganization mechanisms were involved in their response to compression (Figure 6C). The indenter tip was then retracted, and a Hertzian contact model was fitted to the first 10 µm of the unloading curve to determine the reduced elastic modulus (E_r_) (Figure 6B). Although the difference was not statistically significant, biofilms grown on NO_3_^-^medium showed a trend toward higher stiffness, with E_r_ values approximately 55% higher than those of biofilms grown on NH_4_^+^ medium (320 kPa and 180 kPa, respectively), despite the agar stiffness being around 100 kPa for both media (Figure 6D). Regarding the viscoelastic behavior of the biofilms, the effective plasticity showed that half of the energy supplied to the material during the indentation was lost, and biofilms grown on NO_3_^-^ media tended to restore less energy than those grown on NH_4_^+^ media (Figure 6E). Finally, adhesion stress was calculated as the lowest point of the indentation curve over the contact area at the maximum indentation depth (Figure 6F). Biofilms grown on an NO₃⁻ medium tended to be more adhesive (4.2 kPa) than those grown on NH_4_^+^ medium (3.5 kPa).

**Figure 6.**
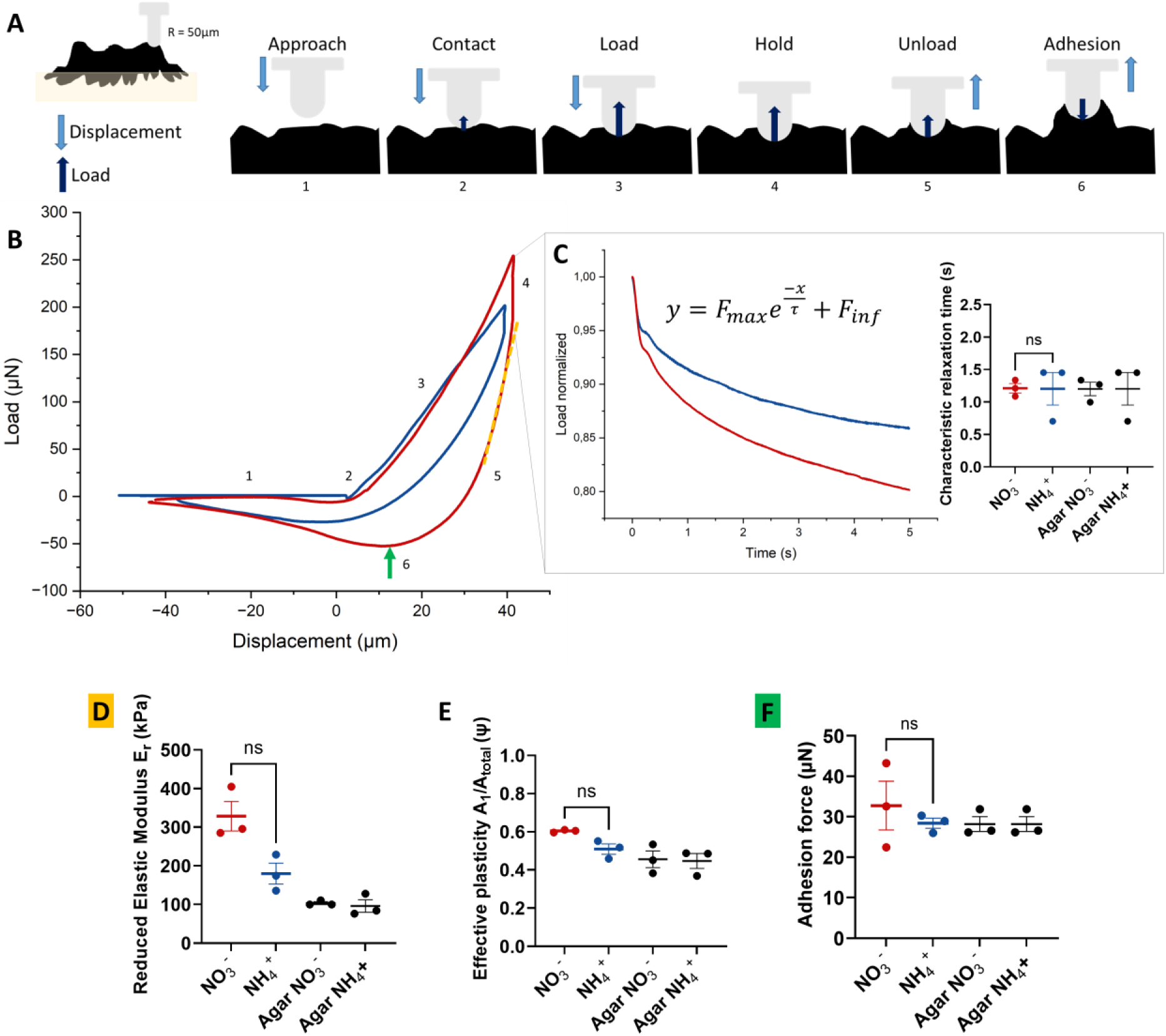
Mechanical characterization of *K. petricola* biofilms through micro-indentation. **(A)** Schematic representation of the experiment. **(B)** Representative load-displacement curves with a holding time obtained on biofilms grown on agar media containing either NO ^-^ (red) or NH ^+^ (blue) as nitrogen source. **(C)** Representative load-time curve describes the relaxation behavior of the material and characteristic relaxation time obtained by fitting an exponential decay function on the load-time curve. **(D)** Reduced elastic moduli (E_r_) obtained on the biofilms and their respective agar substrates fitted with a Hertzian contact function on the 10 first microns of the unloading curve. **(E)** Effective Plasticity is defined as A_1_/A_total_ with A_1_ the area between the loading and unloading curve and A_total_ the area below the loading curve. **(F)** The Adhesion Force was similar for all conditions and samples. Statistical analysis was done with three biological replicates represented with dots on the graphs, and a bar shows the mean value with standard error. A paired two tailed t-test was applied with statistical significance as p < 0.05, * and ns= non-significant.

## Discussion

The mechanical behavior of biological materials is closely linked to their internal structural organization across multiple spatial scales. This relationship has been widely demonstrated in soft biological tissues, where mechanical performance emerges from the hierarchical arrangement of composite structural elements. ^36, 37^ Comparable concepts have also been established for bacterial biofilms, which are increasingly regarded as viscoelastic biological materials or even living fluids, whose macroscopic mechanical properties are determined by cellular packing, EPS composition and architecture, as well as water content. ^6, 38, 39^ In contrast, the biomechanical properties of melanized fungal biofilms and how they integrate in the structure-function relationship of the living material are still relatively poorly resolved.

With this work, we contribute to bridge this gap by establishing a practical experimental workflow that enables structural and mechanical characterization of fungal biofilms across different length scales. The methodological concept was inspired by integrated approaches previously developed in bacterial biofilm research ^5^ but still missing in the current fungal research toolbox. As a model organism, we used the material-inhabiting fungus *K. petricola*, for which growth behavior and morphological transitions are known to be affected by environmental conditions. ^35^ This difference in phenotype at different scales provides a useful framework for exploring how variations in cellular organization translate into differences in emergent biofilm properties. As a first application of this workflow, we compared *K. petricola* biofilms grown under NH_4_^+^ and NO_3_^-^ nitrogen regimes to assess how metabolic conditions influence biofilm structural organization and mechanical behavior.

The methods combined are complementary and altogether provide an integrated view of fungal biofilm architecture. Micro-CT scans enabled visualization of fungal biomass spatial distribution on and in the agar substrate. Three-dimensional reconstructions revealed the overall colony morphology and enabled the quantification of geometrical features such as the volume, surfaces of interactions and thickness distribution (Figure 2). To examine cellular organization and ultrastructure as well as EPS distribution, visualization was complemented with optical imaging of biofilm cryo-sections and with cryo-SEM (Figure 3, Figure 4). Mechanical characterization further benefited from the integration of techniques probing different levels of organization. Shear-rheology describes the bulk viscoelastic behavior of the fungal biomass and thus reports on the collective response of cells and matrix components (Figure 5), whereas micro-indentation reveals local stiffness of the outer layer (Figure 6). These two mechanical tests also differ in their degree of invasiveness and their geometry of mechanical loading. Indeed, rheology implies cutting the biofilms and subjecting the samples to large shear deformations altering their structural integrity, while micro-indentation is performed on native biofilms and localizes compressive load over a few cells and their surrounding EPS, thereby leaving the native structure intact. Considering the important role of water in biofilm structure and mechanical properties, it is important to note that all the techniques used here largely preserved the native hydrated state of the samples.

The methodological approach proposed in this work enables the interpretation of fungal biofilm mechanical behavior considering their hierarchical structure. In the example comparison of *K. petricola* biofilms grown on NH_4_^+^-vs. NO_3_^-^-containing agar substrates, micro-indentation and rheology exhibit consistent trends despite differences in magnitudes due to different probing scales and inherent structural heterogeneities observed in the cross-sections (Figure 4, Figure 5).

Biofilms grown on NO_3_^-^-containing medium are the stiffest and display higher storage and loss moduli than *Candida albicans* BWP17 biofilms, ^40^ whereas NH_4_^+^-biofilms showed lower moduli. *K. petricola* biofilms grown in the presence of NO_3_^-^ exhibited greater similarity to *E. coli* AR3110 bacterial biofilm. ^5^ However, their reduced elastic modulus E_r_ measured by micro-indentation (Figure 6D) is approximately three times higher than the bacterial biofilms. An increased stiffness in the outer layer of *K. petricola* biofilms grown on NO_3_^-^ could be explained by the high concentration of melanized cells observed at the biofilm-air interface (Figure 2). Indeed, melanized cell walls may act as structural reinforcements, significantly enhancing the local mechanical resistance compared to *E. coli* AR3110 embedded in their EPS.^8, 41^

Both mechanical tests showed that biofilms grown on NO_3_^-^-containing medium are stiffer than biofilms grown on NH₄⁺-containing medium, which indicates greater resistance to irreversible deformation (Figure 4C, Figure 6D). This difference can be explained by the cryo-section images revealing NO_3_^-^-cultivated biofilms that are more compact and organized into vertical packages, whereas those grown on NH_4_^+^-containing medium appear less dense and organized in horizontal states (Figure 2A). Drying the cross-sectional slices indicated that these latter also retain more water, especially in the middle layer, which is in line with their reduced water activity (free water) (Figure S4), and likely contributes to their reduced mechanical properties (Figure 4C, Figure 6D). The horizontal and hydrated layering of biofilms cultured on NH_4_^+^-containing medium could also explain their greater tolerance to shear strain (Figure 4E, G). Indeed, the layers may first rather slide with respect to each other before reaching the sudden and irreversible rupture of internal structures (Figure S2). In contrast, the vertical structures observed in biofilms grown on NO_3_^-^ likely undergo a sequential and therefore progressive failure starting at lower shear strains.

While the mechanical characterization methods used here provided insight into the viscoelastic behavior of the emerged part of the biofilm (i.e. above the agar surface), the mechanics of the filamentous network growing within the agar substrate remains not accessible yet. However, micro-CT (Figure 1) and cryo-SEM (Figure 3) imaging revealed distinct structural differences at the interface between the biofilms and their agar substrate, which can help understand their interactions. When NO_3_^-^ is present in the substrate, biofilms penetrate deeper into the substrate by forming millimeter-sized filaments extending from the biofilm edges outwards (Figure 1A) consistent with previous observations. ^35^ Instead, filament penetration in agar supplemented with NH₄⁺ is limited to several hundreds of micrometers, localized in central rings beneath the biofilm and oriented perpendicular to the underlying surface. This divergence may come from multiple factors such as a preference for NH₄⁺, observed for several fungi, ^42^ causing a foraging response with NO_3_^-^, the higher stiffness of NO_3_^-^-grown biofilms, or differences in cell division patterns. For example, the increased mechanical stiffness of biofilms grown on media containing NO_3_^-^ may facilitate their penetration into the agar substrate (Figure 5C, Figure 6D). However, no microscopic deformation of the agar network is visible on cryo-SEM images, which does not support this hypothesis (Figure 4B, C). Instead, the fungal EPS seems to be integrated into the agar network, suggesting that the penetration process occurs during EPS network assembly (e.g., by substituting water in the pores) rather than via mechanical forces. Moreover, cryo-SEM imaging suggested that the nitrogen source modulates the cell-division pattern (Figure 3D, E), where NO_3_^-^ seems to promote budding, whereas NH_4_^+^ seems to favor meristematic division. This shift in developmental strategy may also explain some differences observed in morphology and mechanical properties.

## Conclusion

Taken together, applying the proposed methodological workflow to the present case study revealed that the nitrogen source not only affects *K. petricola* biofilm morphology but also plays a key role in biofilm–substrate interactions as well as in the emergent mechanical properties of the biofilm. Thereby, this work demonstrated the potential of adapting and combining structural and mechanical methods at multiple scales for investigating material-colonizing fungi and addressing a variety of research questions related to fungal biofilm adaptation to genetic modifications and/or environmental growth conditions. The present workflow can be extended by integrating sub-cellular analysis techniques that would enable an additional focus on the matrix components. ^43^ Furthermore, the proposed toolbox offers potential for the systematic investigation of fungal mutant strains, enabling evaluation of how alterations in cell wall composition, such as changes in DHN-melanin deposition, affect biofilm structure and mechanical behavior. Indeed, as is the case in bacterial biofilms, fungal EPS is expected to play a significant role in the functional material properties of fungal biofilms. Understanding this relationship will be beneficial to design innovative strategies to prevent or remove fungal biofilms from substrates, but also to expand the approaches for engineering fungal-based materials.

### Materials and methods Strain and culturing

*K. petricola* strain A95, isolated from a marble monument in Athens, Greece was used in this study ^25^. Prior to the experiments, the strain was cultivated on solid (1.5% bacteriological agar, AppliChem) minimal defined medium containing NO_3_^-^ as the nitrogen source for 7 days at 25°C in darkness. The defined medium contained 100 mM glucose, 10 mM NaNO_3_ or 10 mM NH₄Cl as sole nitrogen sources, and the following salts and trace elements at final concentrations of 5.88 mM KH_2_PO_4_, 6.71 mM KCl, 4.15 mM MgSO_4_, 0.90 mM CaCl_2_, 0.16 µM H_3_BO_3_, 0.025 µM MnCl_2_, 7.55 µM FeSO_4_, 0.80 µM CoCl_2_, 0.10 µM NiCl_2_, 0.059 µM CuCl_2_, 0.50 µM ZnSO_4_, 10 µM NaOH, 0.023 µM Na_2_SeO_3_, and 0.149 µM Na_2_MoO_4_, diluted in ultrapure water (MilliQ). Collected cell aggregates were mechanically dispersed in 500 µL ultrapure water with 4-5 glass beads (3 mm diameter) using a mixer mill (Retsch) at 30 s^-1^ for 5 minutes.

The colony forming units (CFU) of the suspensions were determined using a Thoma counting chamber and adjusted to a final concentration of 10⁷ CFU mL^-1^. Subsequently, 10 µL (10^5^ CFU) was dispensed onto 12.5 mL agar medium containing either NO_3_^-^ or NH_4_^+^ in 6-cm Petri dishes, and incubated for 28 days at 25°C in darkness.

### Micro-Computed Tomography (micro-CT)

To perform a micro-CT with contrast high enough, knowing that agar and biofilm are both hydrogels quite similar in density, iodine staining was applied overnight, according to Metscher ^44^. The biofilms with the agar underneath were placed onto another Petri dish filled with 10% dilution of the stock solution containing 1% iodine metal (I2) and 2% potassium iodine (KI). The solution was removed, and the stained biofilm was placed in a Falcon tube with few more layers of agar underneath to maintain a sufficiently high humidity level.

Micro-CT scans were acquired with an EasyTom Nano 160 system (RX solutions) equipped with a microtube and a horizontal Flat-Panel-Detector. Scanning parameter were set to a voltage of 70 kV, tube current 120 µA and a voxel size of 7.45 µm. The frame rate was set to 1.25 fps, with an average of 10 frames per pass. A total of 1,440 images were taken through four passes to allow for image post-processing, despite the little movement resulting from desiccation during acquisition.

Reconstruction in 3D was done using the XAct software and included spot correction, geometric x-shift correction, and anti-ring correction. Further processing was done with Dragonfly software to segment the biofilm from the agar and obtain data about volume and interfaces. To allow a better contrast and therefore a better segmentation, convolution (in 3D, Kernel size 3, Sphere) and CLAHE (Contrast Limited Adaptative Histogram Equalization) filters were applied before running a deep learning model to identify each zone of the sample. The micro-CT stack of images was therefore segmented in three different parts corresponding to the agar, the biofilm, and the background. This allowed volume measurement of the biofilm but also to compare the interfaces between agar and biofilm and agar and the air (background) with the function “Compute Interfacial measurement” (Figure 2C). The biofilm region of interest (ROI) was transformed into a thickness mesh which provides the histograms of the biofilm thickness calculated as the diameter of a hypothetical sphere that fits within each boundary point (Figure 2B).

### Cryo sectioning and imaging

The biofilm samples were collected with the agar underneath and frozen in an embedding medium for frozen tissue (Leica) to stabilize the biofilm structure. The cryo-microtome from Leica (CM3050) was set at-20°C to cut 30 µm-thick slices out of biofilms on agar or 70 µm-thick slices out of uncolonized agar, all carefully placed on a slide. To image biofilms in a hydrated state, it was covered with a cover slip. However, bubble formation obscured a clear view of the biofilm-agar interaction. Additional images were acquired of agar alone and of agar with biofilm in a dry state, in which the samples were allowed to dry on the glass slide prior to imaging. Imaging was performed with a Zeiss Axiozoom V.16 microscope at 176x digital magnification under brightfield illumination, using the automatic white balance correction.

### Cryogenic Scanning Electron Microscopy (Cryo-SEM)

Cryo-SEM was performed with a Zeiss Crossbeam 550 FIB/SEM equipped with a Leica cryo transfer system. A piece of agar and biofilm were cut and placed on a 2 mm-diameter sample holder, with the cross section facing upward. The specimens were then plunged into a nitrogen slush for a few seconds. The frozen specimens were vacuum transferred at cryo temperature to an ACE900 freeze fracture system using a VCT500 cryo transfer system. Freeze fracture was performed with a stage temperature of-170°C, followed by a freeze-etching at 100°C for 5 minutes to expose the internal pore structures. A 5nm-thick platinum layer was e-beam deposited on the specimens at-170°C to enhance electrical conductivity of the specimen surface and better protection against electron radiation damage. The specimens were transferred to the cryo-FIB/SEM and imaged at a stage temperature of-160°C using In-lens secondary electron and In-chamber secondary electron/secondary ion detectors.

### Rheology

The viscoelastic properties of the fungal biofilms were analyzed using a rheometer (MCR301, Anton Paar GmbH). About one quarter of the biofilm was carefully excised and placed on the rheometer plate to avoid structural disruption. Measurements were performed at 21°C (Stage Peltier thermoelectric cooling) with a cone plate geometry of 12 mm diameter, and a gap set at ∼100 µm. The measurement setup was enclosed with a hood to prevent sample dehydration by maintaining humidity.

One frequency sweep was performed on half of the biofilm at 1% strain and oscillating between 0.01 rad/s to 1000 rad/s for each condition. The frequency was then set at 10 rad/s, and the other half of the biofilm as well as the other biological replicates were subjected to amplitude sweeps from 0.01 to 1000% of deformation with 7 points per decade.

The linear viscoelastic region (LVR) was identified from strain-amplitude sweep experiments as the plateau region of the storage modulus (G′), over which both G′ and the loss modulus (G″) remained approximately independent of strain amplitude. The storage modulus (G′) and loss modulus (G″) were recorded as functions of applied strain. The plateau storage modulus G′_0_ was determined as the mean value of G′ within the LVR, and the LVR limit was defined as the strain at which G′ decreased to 95% of G′_0_. This criterion was used to characterize the elastic response of the biofilm. For each applied strain amplitude γ, the magnitude of the complex shear modulus G was calculated as the square root of the sum of the squared storage modulus (G′) and squared loss modulus (G″). The corresponding shear stress amplitude τ was then obtained by multiplying G by the applied shear strain γ. Strain is reported as γ (dimensionless; values given in percent were converted to fractional form for calculations), and shear stress is reported as τ in pascals (Pa). The transition index (TI) was calculated to quantify the breadth of the yielding-to-flow transition. Specifically, TI was defined as the ratio of the shear stress at the flow point (τ_flow_) to the shear stress at the yield point (τ_yield_). Rheological data were analyzed using R.

### Micro-indentation

Micro-indentation was performed on native biofilms on the agar substrate prior to destructive rheological measurements on the same samples. The TI 950 Triboindenter (Hysitron) was equipped with an extended displacement stage (XZ-500) that allowed for a maximum displacement of 500 µm in the indentation direction and a conospherical tip with a diameter of 50 µm. Calibration was performed in air.

Two configurations were used to obtain load-displacement curves: one with a loading and an unloading phase and another with an additional 5-second holding time at the maximum indentation depth to assess the biofilm’s relaxation behavior. For each configuration and biofilm measured (nine samples: three biological and three technical replicates), two measurements were acquired on the agar, and 4 to 6 measurements were acquired at random locations on the biofilm.

A MATLAB code was used to analyze these data. The indentation starting point was defined as a drastic change in slope and set as the derivative of force over displacement, which equaled 1 for NO₃⁻ and 1.5 for NH₄⁺ and agar. To determine the reduced elastic modulus (E_r_), a Hertzian contact model (see Equation 1) was fitted to the first 25 microns of indentation during loading and the first 10 microns of retraction during unloading.

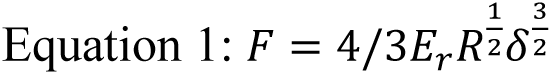

where F is the Force in N, E_r_ is the reduced elastic modulus in Pa, R is the tip radius in m and δ is the displacement in m.

The end of the loading phase (maximum force and displacement) was chosen as the starting point of the time-force relaxation curve for a measurement with a holding time. Information about the relaxation behavior of material was extracted using the characteristic relaxation time of the material obtained from an exponential decay fitted to the relaxation curve (see Equation 2).

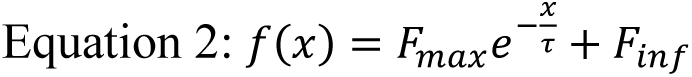

where F_max_ is the maximum force, τ is the characteristic time, and F_inf_ is the remaining force after infinite relaxation.

Relative plasticity refers to how much the material can reorganize around the tip. These indexes are calculated by using the ratio between A_1_ (the area between loading and unloading curve), or A_h_ (the area between loading and unloading curve but only for the holding time) and A_total_ which correspond to the entire area under the loading curve.

The curve minimum after retraction was set as the adhesion force of the material.

### Water activity

Water activity of the samples was determined using a water activity meter (LabMaster-aw Neo, Novasina AG). Measurements were performed on three distinct sample types: [i] the biofilm, [ii] the agar underneath the biofilm containing some filaments, and [iii] the agar medium. For each condition, approximately 1 cm^2^ of material was carefully collected to avoid moisture loss. Samples were placed in the measurement chamber immediately after sampling and equilibrated according to the instrument protocol until a stable reading was obtained. The device was calibrated prior to each measurement session using standard salt solutions of known water activity values (97%). Osmotic pressure (Π) was calculated using the thermodynamic relationship derived from van’t Hoff’s law, which relates the water activity of a solution to its osmotic pressure. Specifically, 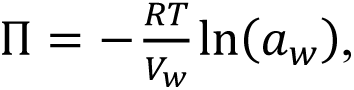 where V_w_ is the molar volume of water in solution (V_w_ = 1.8 × 10^-5^ m^3^/mol), R is the universal gas constant (R = 8.314 m^3^.Pa/(K.mol)), and T is the absolute temperature (T = 298.15 K). For each sample, water activity values were used in this equation to compute osmotic pressure. The osmotic gradient (ΔΠ) between the biofilm and agar was further defined as: 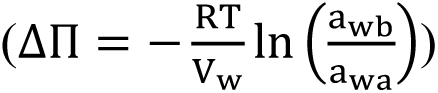

## Statistical analysis

The statistics for the mechanical experiments were performed using GraphPad Prism and Origin software. Three biological replicates (grown from different precultures and inoculated on different days) and three technical replicates of NO_3_^-^ and NH_4_^+^ were measured. For each biological replicate, inoculations for the two tested conditions were done on the same day with the same cultures to avoid influence from environmental conditions; therefore, our data was considered dependent. The normality of the data was tested with the Shapiro-Wilk method and analyzed with a quantile test plot. A parametric test was chosen because the data was considered normally distributed. Therefore, a paired two-sided t-test was performed, and the mean of difference can be found in Figure S3.

## Contributions

Conceptualization: C.F., A.D., J.S., R.G., P.F., A.A.G., C.M.B.

Experimental developments: C.F., A.D., Y.O.

Data acquisition and analysis: C.F., A.D., Y.O., C.M.B. Manuscript preparation and writing: C.F., A.D., J.S., Y.O., C.M.B. Manuscript reviewing and editing: all co-authors

## Acknowledgments

The authors thank Christine Pilz-Allen, Susann Weichold, Daniel Werner, Shahrouz Amini and Nives Hribernik for their technical support in the laboratories. They also thank Laura Zorzetto, Emeline Raguin and Angelo Valleriani for their help in analyzing data.

This article is based upon work from COST Action EuroCurvoBioNet CA22153, supported by COST (European Cooperation in Science and Technology).

## Supporting Information

Complementary data for micro-CT analysis, Rheological characterization of fungal biofilms, Statistical analysis of rheology and micro-indentation data, Water activity of biofilms, agar underneath biofilms and agar prepared with different nitrogen sources.

## References

(1) Flemming, H.-C.; van Hullebusch, E. D.; Neu, T. R.; Nielsen, P. H.; Seviour, T.; Stoodley, P.; Wingender, J.; Wuertz, S. The biofilm matrix: multitasking in a shared space. Nat. Rev. Microbiol. 2023, 21 (2), 70–86, DOI: 10.1038/s41579-022-00791-0.

(2) Ziege, R.; Tsirigoni, A.-M.; Large, B.; Serra, D. O.; Blank, K. G.; Hengge, R.; Fratzl, P.; Bidan, C. M. Adaptation of *Escherichia coli* biofilm growth, morphology, and mechanical properties to substrate water content. ACS Biomater. Sci. Eng. 2021, 7 (11), 5315–5325, DOI: 10.1021/acsbiomaterials.1c00927.

(3) Tallawi, M.; Opitz, M.; Lieleg, O. Modulation of the mechanical properties of bacterial biofilms in response to environmental challenges. Biomater. Sci. 2017, 5 (5), 887–900, DOI: 10.1039/c6bm00832a.

(4) Siri, M.; Vázquez-Dávila, M.; Sotelo Guzman, C.; Bidan, C. M. Nutrient availability influences *E. coli* biofilm properties and the structure of purified curli amyloid fibers. npj Biofilms Microbiomes 2024, 10 (1), 143, DOI: 10.1038/s41522-024-00619-0.

(5) Siri, M.; Sarlet, A.; Ziege, R.; Zorzetto, L.; Sotelo Guzman, C.; Amini, S.; Hengge, R.; Blank, K. G.; Bidan, C. M. Mechanical comparison of *Escherichia coli* biofilms with altered matrix composition: a study combining shear-rheology and microindentation. ACS Biomater. Sci. Eng. 2025, 11 (7), 4523–4536, DOI: 10.1021/acsbiomaterials.5c00261.

(6) Boudarel, H.; Mathias, J.-D.; Blaysat, B.; Grédiac, M. Towards standardized mechanical characterization of microbial biofilms: analysis and critical review. npj Biofilms Microbiomes 2018, 4 (1), 17, DOI: 10.1038/s41522-018-0062-5.

(7) Serra, D. O.; Richter, A. M.; Klauck, G.; Mika, F.; Hengge, R. Microanatomy at cellular resolution and spatial order of physiological differentiation in a bacterial biofilm. mBio 2013, 4 (2),doi:10.1128/mbio.00103-13.

(8) Serra, D. O.; Richter, A. M.; Hengge, R. Cellulose as an architectural element in spatially structured *Escherichia coli* biofilms. J. Bacteriol. 2013, 195 (24), 5540–5554, DOI: 10.1128/jb.00946-13.

(9) Schröer, L.; Balcaen, T.; Folens, K.; Boon, N.; De Kock, T.; Kerckhofs, G.; Cnudde, V. 3D Visualization of cyanobacterial biofilms using micro-computed tomography with contrast-enhancing staining agents. *Tomogr*. Mater. Struct. 2024, 4, 100024, DOI: 10.1016/j.tmater.2024.100024.

(10) Lieleg, O.; Caldara, M.; Baumgärtel, R.; Ribbeck, K. Mechanical robustness of *Pseudomonas aeruginosa* biofilms. Soft Matter 2011, 7 (7), 3307–3314, DOI: 10.1039/C0SM01467B.

(11) Liu, S.; Huang, J.; Zhang, C.; Wang, L.; Fan, C.; Zhong, C. Probing the growth and mechanical properties of *Bacillus subtilis* biofilms through genetic mutation strategies. Synth. Syst. Biotechnol. 2022, 7 (3), 965–971, DOI: 10.1016/j.synbio.2022.05.005.

(12) Siri, M.; Mangiarotti, A.; Seewald, A.; Rosenthal, N.; Amini, S.; Raguin, E.; Fratzl, P.; Bidan, C. M. *E. coli* extracellular matrix: a tunable composite with hierarchical structure. bioRxiv 2026, 1–35, DOI: 10.64898/2026.02.22.707275.

(13) Blankenship, J. R.; Mitchell, A. P. How to build a biofilm: a fungal perspective. Curr Opin Microbiol 2006, 9 (6), 588–594, DOI: 10.1016/j.mib.2006.10.003.

(14) Ramage, G.; Mowat, E.; Jones, B.; Williams, C.; Lopez-Ribot, J. Our current understanding of fungal biofilms. Crit Rev Microbiol 2009, 35 (4), 340–355, DOI: 10.3109/10408410903241436.

(15) Gow, N. A. R.; Latge, J.-P.; Munro, C. A. The fungal cell wall: structure, biosynthesis, and function. Microbiol. Spectr. 2017, 5 (3), 1–25, DOI: 10.1128/microbiolspec.funk-0035-2016.

(16) Breitenbach, R.; Toepel, J.; Dementyeva, P.; Knabe, N.; Gorbushina, A. Snapshots of fungal extracellular matrices. In *The Perfect Slime: Microbial Extracellular Polymeric Substances (EPS)*, Flemming, H.-C., Neu, T. R., Wingender, J. Eds.; London: IWA Publishing, 2016; pp 269–299, DOI: 10.2166/9781780407425.

(17) Sudbery, P.; Gow, N.; Berman, J. The distinct morphogenic states of *Candida albicans*. Trends Microbiol. 2004, 12 (7), 317–324, DOI: 10.1016/j.tim.2004.05.008.

(18) Mitchison-Field, L. M. Y.; Vargas-Muñiz, J. M.; Stormo, B. M.; Vogt, E. J. D.; Van Dierdonck, S.; Pelletier, J. F.; Ehrlich, C.; Lew, D. J.; Field, C. M.; Gladfelter, A. S. Unconventional cell division cycles from marine-derived yeasts. Curr. Biol. 2019, 29 (20), 3439–3456.e3435, DOI: 10.1016/j.cub.2019.08.050.

(19) Wollenzien, U.; De Hoog, G.; Krumbein, W.; Urzi, C. On the isolation of microcolonial fungi occurring on and in marble and other calcareous rocks. Sci Total Environ 1995, 167 (1-3), 287–294, DOI: 10.1016/0048-9697(95)04589-S.

(20) De Leo, F.; Marchetta, A.; Urzì, C. Black fungi on stone-built heritage: current knowledge and future outlook. Appl. Sci. 2022, 12 (8), 3969, DOI: 10.3390/app12083969.

(21) Tesei, D. Black fungi research: out-of-this-world implications. Encyclopedia 2022, 2 (1), 212–229, DOI: 10.3390/encyclopedia2010013.

(22) Gorbushina, A. A.; Krumbein, W. E.; Hamman, C. H.; Panina, L.; Soukharjevski, S.; Wollenzien, U. Role of black fungi in color change and biodeterioration of antique marbles. Geomicrobiol J 1993, 11 (3-4), 205–221, DOI: 10.1080/01490459309377952.

(23) Martin-Sanchez, P. M.; Gebhardt, C.; Toepel, J.; Barry, J.; Munzke, N.; Gunster, J.; Gorbushina, A. A. Monitoring microbial soiling in photovoltaic systems: A qPCR-based approach. Int Biodeter Biodegr 2018, 129, 13–22, DOI: 10.1016/j.ibiod.2017.12.008.

(24) Noack-Schönmann, S.; Spagin, O.; Gründer, K. P.; Breithaupt, M.; Günter, A.; Muschik, B.; Gorbushina, A. A. Sub-aerial biofilms as blockers of solar radiation: spectral properties as tools to characterise material-relevant microbial growth. Int Biodeter Biodegr 2014, 86, 286–293, DOI: 10.1016/j.ibiod.2013.09.020.

(25) Gorbushina, A. A.; Kotlova, E. R.; Sherstneva, O. A. Cellular responses of microcolonial rock fungi to long-term desiccation and subsequent rehydration. Stud. Mycol. 2008, 61, 91–97, DOI: 10.3114/sim.2008.61.09.

(26) Nai, C.; Wong, H. Y.; Pannenbecker, A.; Broughton, W. J.; Benoit, I.; de Vries, R. P.; Gueidan, C.; Gorbushina, A. A. Nutritional physiology of a rock-inhabiting, model microcolonial fungus from an ancestral lineage of the Chaetothyriales (Ascomycetes). Fungal Genet. Biol. 2013, 56, 54–66, DOI: 10.1016/j.fgb.2013.04.001.

(27) Wollenzien, U.; De Hoog, G.; Krumbein, W.; Uijthof, J. *Sarcinomyces petricola*, a new microcolonial fungus from marble in the Mediterranean basin. Antonie van Leeuwenhoek 1997, 71 (3), 281–288, DOI: 10.1023/a:1000157803954.

(28) Gorbushina, A. A.; Broughton, W. J. Microbiology of the atmosphere-rock interface: how biological interactions and physical stresses modulate a sophisticated microbial ecosystem. Annu Rev Microbiol 2009, 63, 431–450, DOI: 10.1146/annurev.micro.091208.073349.

(29) Breitenbach, R.; Gerrits, R.; Dementyeva, P.; Knabe, N.; Schumacher, J.; Feldmann, I.; Radnik, J.; Ryo, M.; Gorbushina, A. A. The role of extracellular polymeric substances of fungal biofilms in mineral attachment and weathering. *npj Mater*. Degrad. 2022, 6 (1), 42, DOI: 10.1038/s41529-022-00253-1.

(30) Gerrits, R.; Wirth, R.; Schreiber, A.; Feldmann, I.; Knabe, N.; Schott, J.; Benning, L. G.; Gorbushina, A. A. High-resolution imaging of fungal biofilm-induced olivine weathering. Chemical. Geol. 2021, 559, DOI: 10.1016/j.chemgeo.2020.119902.

(31) Gerrits, R.; Pokharel, R.; Breitenbach, R.; Radnik, J.; Feldmann, I.; Schuessler, J. A.; von Blanckenburg, F.; Gorbushina, A. A.; Schott, J. How the rock-inhabiting fungus *K. petricola* A95 enhances olivine dissolution through attachment. Geochim. Cosmochim. Ac. 2020, 282, 76–97, DOI: 10.1016/j.gca.2020.05.010.

(32) Breitenbach, R.; Silbernagl, D.; Toepel, J.; Sturm, H.; Broughton, W. J.; Sassaki, G. L.; Gorbushina, A. A. Corrosive extracellular polysaccharides of the rock-inhabiting model fungus *Knufia petricola*. Extremophiles 2018, 22 (2), 165–175, DOI: 10.1007/s00792-017-0984-5.

(33) Voigt, O.; Knabe, N.; Nitsche, S.; Erdmann, E. A.; Schumacher, J.; Gorbushina, A. A. An advanced genetic toolkit for exploring the biology of the rock-inhabiting black fungus *Knufia petricola*. Sci. Rep. 2020, 10 (1), 22021, DOI: 10.1038/s41598-020-79120-5.

(34) Flieger, K.; Knabe, N.; Toepel, J. Development of an improved carotenoid extraction method to characterize the carotenoid composition under oxidative stress and cold temperature in the rock inhabiting fungus *Knufia petricola* A95. J Fungi 2018, 4 (4), 124, DOI: 10.3390/jof4040124.

(35) Dehkohneh, A.; Schumacher, J.; Cockx, B. J. R.; Keil, K.; Camenzind, T.; Kreft, J.-U.; Gorbushina, A. A.; Gerrits, R. Carbon and nitrogen availability affect biofilm growth and morphology of the extremotolerant fungus *Knufia petricola*. bioRxiv 2026, 1–35, DOI: 10.64898/2026.03.19.712823.

(36) Dunlop, J. W. C.; Fratzl, P. Multilevel architectures in natural materials. Scripta Mater 2013, 68 (1), 8–12, DOI: 10.1016/j.scriptamat.2012.05.045.

(37) Fratzl, P. Hierarchical structure and mechanical adaptation of biological materials. Dordrecht, 2005; Springer Netherlands: pp 15–34. DOI: 10.1007/1-4020-2648-X_2.

(38) Wells, M. J.; Zhou, X.; Gordon, V. D. Viscoelasticity of the Biofilm Matrix. In Biofilm Matrix, Reichhardt, C. Ed.; Springer Nature Switzerland, 2024; pp 259–282, DOI: 10.1007/978-3-031-70476-5_8.

(39) Wilking, J. N.; Angelini, T. E.; Seminara, A.; Brenner, M. P.; Weitz, D. A. Biofilms as complex fluids. MRS Bulletin 2011, 36 (5), 385–391, DOI: 10.1557/mrs.2011.71.

(40) Beckwith, J. K.; Ganesan, M.; VanEpps, J. S.; Kumar, A.; Solomon, M. J. Rheology of *Candida albicans* fungal biofilms. J. Rheol. 2022, 66 (4), 683–697, DOI: 10.1122/8.0000427.

(41) Mattoon, E. R.; Cordero, R. J. B.; Casadevall, A. Melaninization reduces *Cryptococcus neoformans* susceptibility to mechanical stress. mSphere 2023, 8 (1), e00591–00522,doi:10.1128/msphere.00591-22.

(42) Finlay, R. D.; FrostegÅRd, Å.; Sonnerfeldt, A. M. Utilization of organic and inorganic nitrogen sources by ectomycorrhizal fungi in pure culture and in symbiosis with Pinus contorta Dougl. ex Loud. New Phytologist 1992, 120 (1), 105–115, DOI: 10.1111/j.1469-8137.1992.tb01063.x.

(43) Pham, D. Q.; Bryant, S. J.; Cheeseman, S.; Huang, L. Z. Y.; Bryant, G.; Dupont, M. F.; Chapman, J.; Berndt, C. C.; Vongsvivut, J.; Crawford, R. J.; Truong, V. K.; Ang, A. S. M.; Elbourne, A. Micro-to nano-scale chemical and mechanical mapping of antimicrobial-resistant fungal biofilms. Nanoscale 2020, 12 (38), 19888–19904, DOI: 10.1039/D0NR05617K.

(44) Metscher, B. D. MicroCT for comparative morphology: simple staining methods allow high-contrast 3D imaging of diverse non-mineralized animal tissues. BMC Physiol. 2009, 9 (1), 11, DOI: 10.1186/1472-6793-9-11.

