## Supplementary information for "Biophysical properties of material-colonizing fungal biofilms: multiscale structural and mechanical characterization"

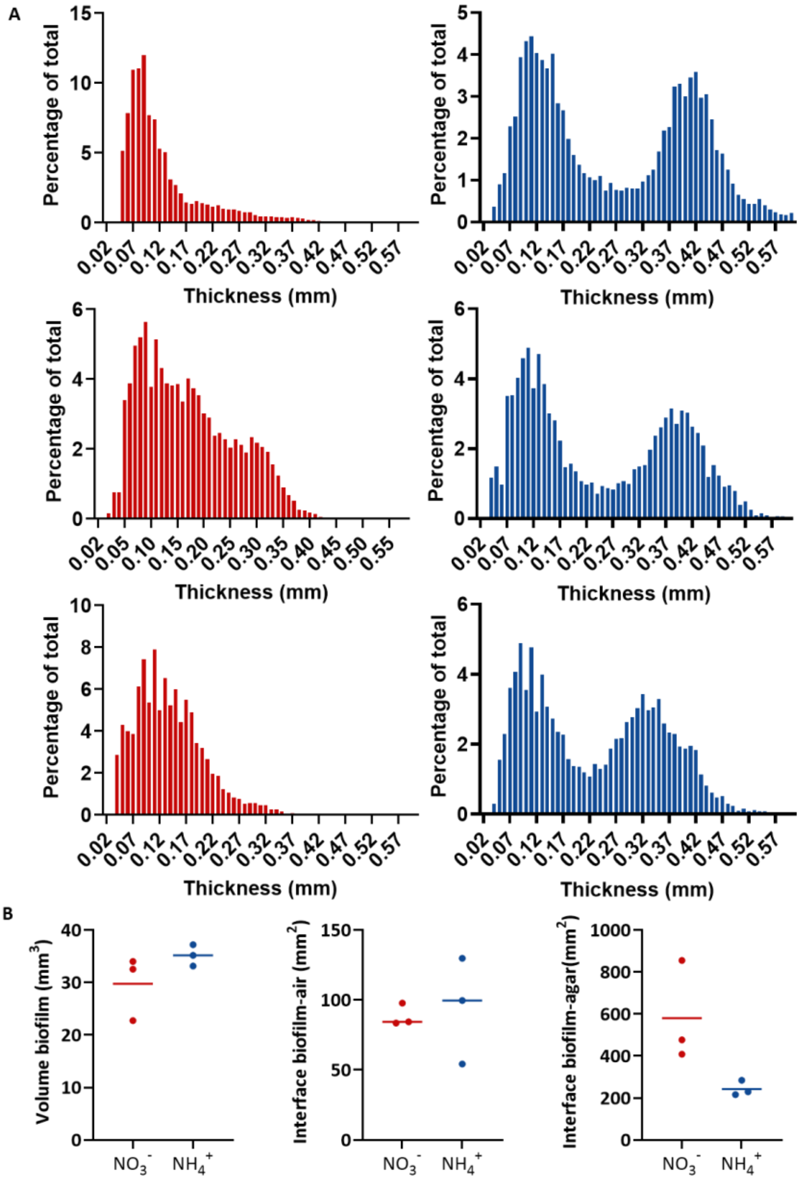


**Figure S1. Complementary data for micro-CT analysis. (A)** Thickness distribution of biofilms for each biological replicate. **(B)** Volume and interfaces calculated with Dragonfly software.

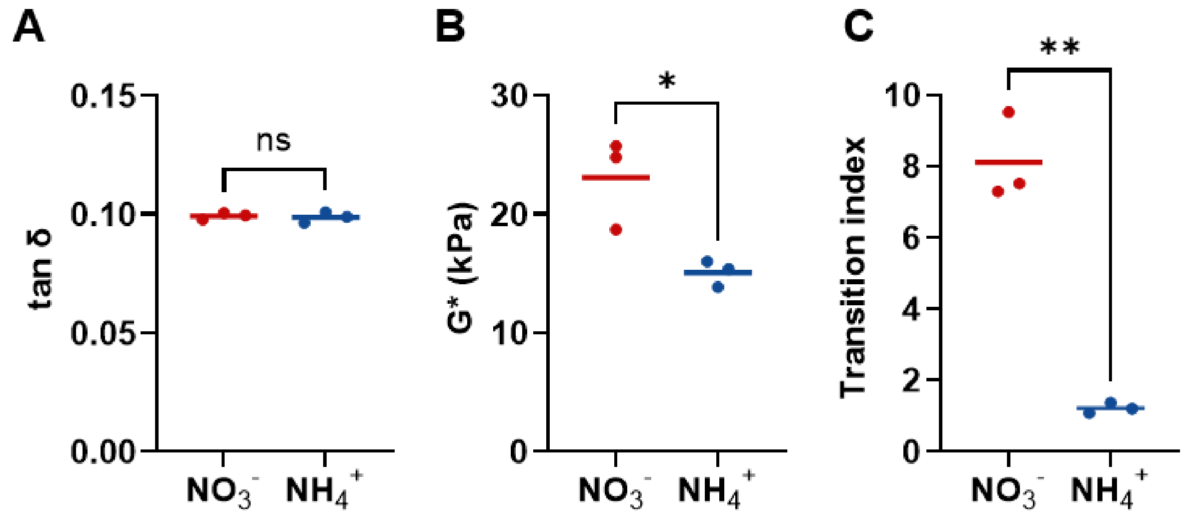


**Figure S2. Rheological characterization of fungal biofilms. (A)** Loss tangent (tan γ). **(B)** Complex modulus (G*). **(C)** Transition index describing the relative dominance of viscous versus elastic behavior (TI = $\tau_{flow}$/ $\tau_{yield}$). Measurements were performed under oscillatory shear, and values represent independent replicates.

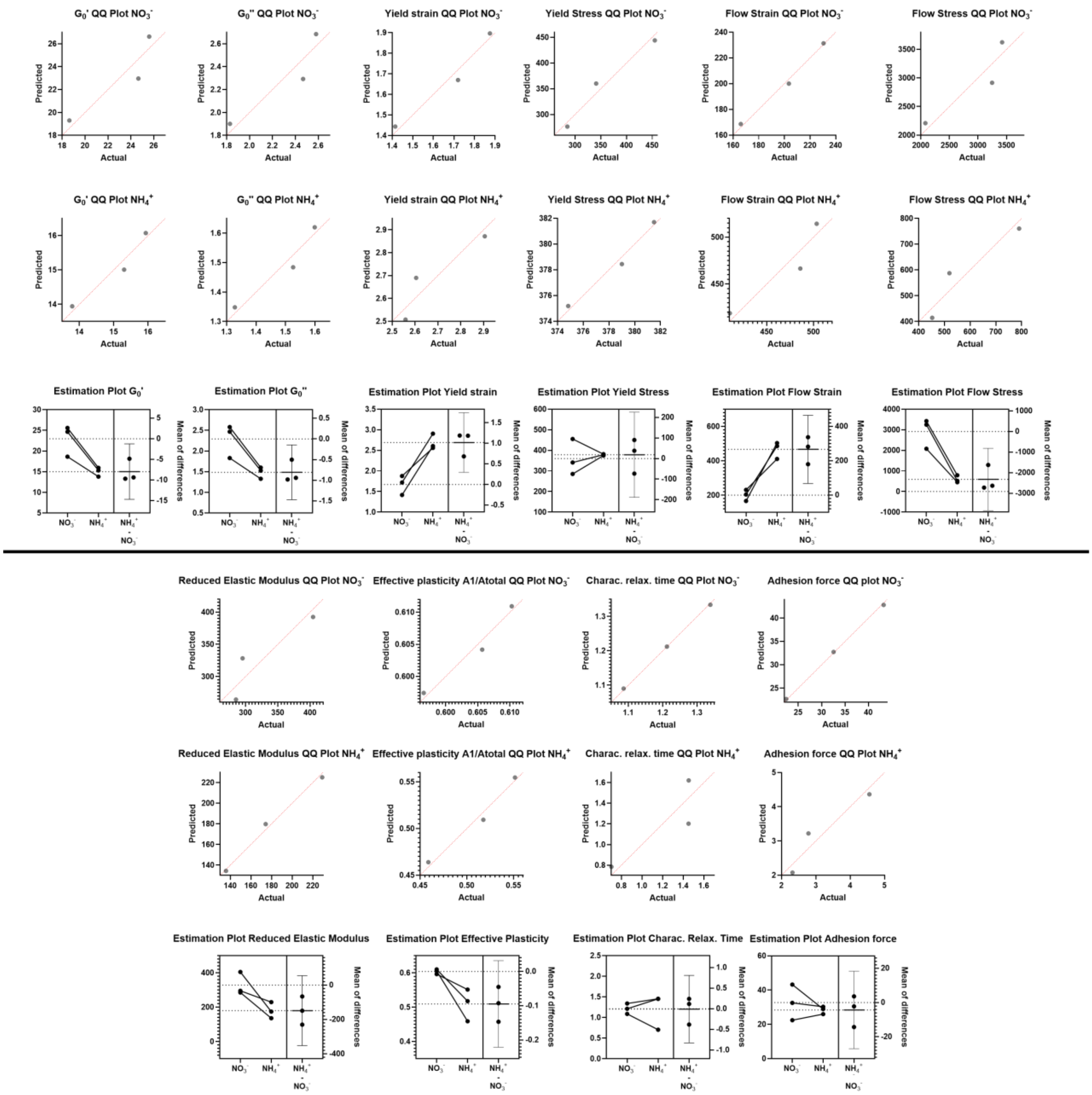


**Figure S3. Statistical analysis of rheology and micro-indentation data.** QQ plots showing data normality distribution for each measured conditions and estimation plots showing data pairing and the mean of difference. All dots represent biological replicates.

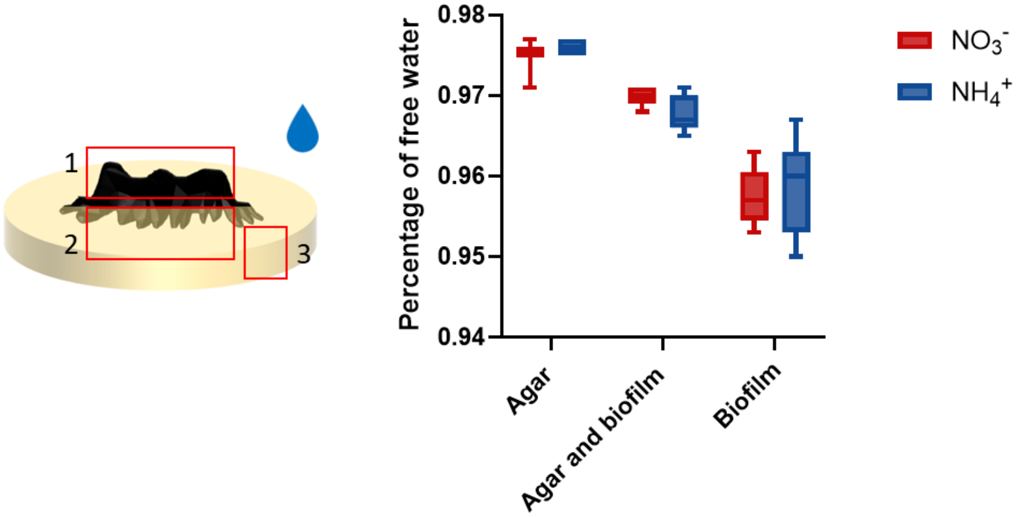


**Figure S4. Water activity of biofilms (1), agar underneath biofilms (2) and agar (3) prepared with different nitrogen sources.** Independent two-sample t-tests comparing and groups revealed non-significant effects of nitrogen source on biofilms, agar underneath of biofilm, and agar measurements (all $p > 0.05$). All experiments were conducted in triplicate.
